# Deep Convolutional Neural Networks Enable Discrimination of Heterogeneous Digital Pathology Images

**DOI:** 10.1101/197517

**Authors:** Pegah Khosravi, Ehsan Kazemi, Marcin Imielinski, Olivier Elemento, Iman Hajirasouliha

**Affiliations:** Institute for Computational Biomedicine, Weill Cornell Medical College, NY, USA; Department of Physiology and Biophysics, Weill Cornell Medicine, New York, NY, USA; Yale Institute for Network Science, Yale University, New Haven, CT, USA; Englander Institute for Precision Medicine, Weill Cornell Medical College, NY, USA; Department of Pathology and Laboratory Medicine, Weill Cornell Medical College, NY, USA; The New York Genome Center, NY, USA; The Meyer Cancer Center, Weill Cornell Medicine, New York, NY, USA

**Author notes:** Contributed equally (Pegah Khosravi, Ehsan Kazemi).

## Abstract

Pathological evaluation of tumor tissue is pivotal for diagnosis in cancer patients and automated image analysis approaches have great potential to increase precision of diagnosis and help reduce human error.

In this study, we utilize various computational methods based on convolutional neural networks (CNN) and build a stand-alone pipeline to effectively classify different histopathology images across different types of cancer. In particular, we demonstrate the utility of our pipeline to discriminate between two subtypes of lung cancer, four biomarkers of bladder cancer, and five biomarkers of breast cancer. In addition, we apply our pipeline to discriminate among four immunohistochemistry (IHC) staining scores of bladder and breast cancers.

Our classification pipeline utilizes a basic architecture of CNN, Google’s Inceptions within three training strategies, and an ensemble of two state-of-the-art algorithms, Inception and ResNet. These strategies include training the last layer of Google’s Inceptions, training the network from scratch, and fine-tunning the parameters for our data using two pre-trained version of Google’s Inception architectures, Inception-V1 and Inception-V3.

We demonstrate the power of deep learning approaches for identifying cancer subtypes, and the robustness of Google’s Inceptions even in presence of extensive tumor heterogeneity. Our pipeline on average achieved accuracies of 100%, 92%, 95%, and 69% for discrimination of various cancer types, subtypes, biomarkers, and scores, respectively. Our pipeline and related documentation is freely available at https://github.com/ih-lab/CNN_Smoothie.

## 1 Introduction

Evaluation of microscopic histopathology slides by experienced pathologists is currently the standard procedure for establishing a diagnosis and identifying the subtypes of different cancers. Visual-only assessment of well-established histopathology patterns is typically slow, and is shown to be inaccurate and irreproducible in certain diagnosis cases of tumor subtypes and stages [58]. Several recent studies attempted to employ machine learning approaches for determining subtypes of malignancies [19, 87]. These computational approaches can be complementary with other clinical evaluation methods to improve pathologists’ knowledge of the disease and improve treatments [21, 4]. For example, previous studies have shown more accurate diagnosis results are derived by integrating information extracted from computational pathology with patients’ clinical data for various cancer types such as prostate cancer [6, 17], lung cancer [28], breast cancer [83, 16], colorectal cancer [42], and ovarian cancer [36]. In particular, computerized image processing technology has been shown to improve performance, correctness, and robustness in histopathology assessments [47].

While new advanced approaches have improved image recognition (e.g., normal versus cancerous), the image interpretation of heterogeneous populations still suffers from lack of robust computerization approaches [66, 11, 26, 37]. Current available automatic methods focus on classification of just one type of cancer versus the corresponding normal condition. Although these studies achieved reasonable accuracy in detecting normal or cancerous conditions in specific kind of cancers, leveraging methods such as training Convolutional Neural Networks (CNNs)[46], they have certain limitations which we address in this work:

1. Developing *ensemble* deep learning methods to employ state-of-the-art algorithms for improving training approaches in diagnosis and detection of various cancer subtypes (e.g., adenocarcinoma versus cell squamous lung cancer).
2. Improving the speed of deep learning, and investigating the trade-offs between performance (i.e., the size of the training set) and efficiency (i.e., the training speed).
3. Making decisions on selecting proper neural networks for different types of data set.

One of the main challenges of computational pathology is that tumor tissue images often vary in color and scale batch effects across different research laboratories and medical facilities due to differences in tissue preparation methods and imaging implements [43]. Previous studies have shown that technicians’ variance or technique differences lead to differences in staining substantially [55] also causes difficulties in extracting clinical information robustly. Furthermore, erroneous evaluation of histopathology images and decision-making using tissue slides containing millions of cells can be time-consuming and subjective [87, 43].

In addition, cancer is known to be a heterogeneous disease. i.e., a high degree of genetic and phenotypic diversity exists “within tumors” (intra-tumor) and/or “among tumors” (inter-tumor) [64]. Tumor heterogeneity leads to an important effect of disease progression and resistant responses to targeted therapies [30]. We also aim to evaluate deep learning approaches for discrimination of digital pathology images from intra- and inter-tumor heterogeneous samples.

Deep learning approaches are emerging as leading machine-learning tools in medical imaging where they have been proven to produce precious results on various tasks such as segmentation, classification, and prediction[24]. In this paper, we present an innovative deep learning based pipeline, CNN_Smoothie, to discriminate various cancer types, subtypes, and their relative staining markers and scores.We combine pathological images of three cancer types with the ones related to the immunohistochemical markers of tumor differentiation to train CNNs for analyzing and identifying specific clinical patterns in different staining markers and scores of breast and bladder cancers. In addition, we applied deep learning methods on immunohistochemistry (IHC) and hematoxylin & esoin (H&E) stained images of squamous cell carcinoma and lung adenocarcinoma to investigate the performance of various classifiers.

To the best of our knowledge, this is the first comprehensive study of applying a wide range of CNN architectures (all integrated in a single pipeline) on histopathology images from multiple different datasets. We evaluate performance of different architectures to detect and diagnosis of tumor images. Our results clearly demonstrate the power of deep learning approaches for distinguishing different cancer types, subtypes, IHC markers and their expression scores. Source codes and documentation of our pipeline containing training, evaluation and prediction methods are publicly available at https://github.com/ih-lab/CNN_Smoothie.

## 2 Materials and Methods

### 2.1 Histopathology images resource

Our datasets come from a combination of open-access histopathology images, The Stanford Tissue Microarray Database (TMAD) and The Cancer Genome Atlas (TCGA). A total of 12139 whole-slide stained histopathology images were obtained from TMAD [53]. TMA database enables researchers have access to bright field and fluorescence images of tissue microarrays. This archive provide thousand human tissues which are probed by antibodies simultaneously for detection of protein abundance (immunohistochemistry; IHC), or by labeled nucleic acids (in situ hybridization; ISH) to detect transcript abundance. The extracted data included samples from three cancer types: (1) lung, (2) breast, comprising five biomarker types (EGFR, CK17, CK5/6, ER, and HER2), and (3) bladder with four biomarker types (CK14, GATA3, S0084, and S100P). Characteristics of all three cohorts and the comprised classes are summarized in Table 1. From the extracted TMA datasets, one dataset is stained by H&E method (BladderBreastLung) and one dataset is stained by both H&E and IHC methods (TMAD-InterHeterogeneity). The remaining datasets (BladderBiomarkers, BreastBiomarkers, BladderScores, and BreastScores) are stained by IHC markers including different polyclonal antiserums such as CK14, GATA3, S0084, S100P, EGFR, CK17, CK5/6, ER, and HER2 for their related proteins which play critical roles in tumor progression.

**Table 1:** Eight datasets are selected to assess the performance of the pipeline across different conditions.

| Number | Datasets | The database representation | Labels of inputs and outputs | Dataset size | Class size |
| --- | --- | --- | --- | --- | --- |
| 1 | BladderBreastLung | H&E-stained images for bladder, breast and lung cancers | Discrimination of different types of cancer (bladder, breast, and lung) | 3 classes and 1918 images | bladder: 543, breast: 962, lung: 413 |
| 2 | BladderBiomarkers | IHC-stained images of cancer biomarkers comprising GATA3, CK14, S100P, and S0084 in bladder cancer | Discrimination of different types of biomarkers (GATA3, CK14, S100P, and S0084) | 4 classes and 2139 images | GATA3: 542, CK14: 514, S100P: 544, S0084: 539 |
| 3 | BreastBiomarkers | IHC-stained images of cancer biomarkers including ER, CK17, CK5/6, EGFR, and HER2 in breast cancer | Discrimination of different types of biomarkers (ER, CK17, CK5/6, EGFR, and HER2) | 5 classes and 2542 images | ER: 637, CK17: 639, CK5/6: 635, EGFR: 307, HER2: 324 |
| 4 | TMAD-InterHeterogeneity | H&E- and IHC-stained whole-slides of adenocarcinoma and squamous cell lung cancers | Discrimination of different subtypes of cancer (adenocarcinoma vs. squamous cell lung tumors) for TMAD images | 2 classes and 860 images (H&E: 572, IHC: 288) | adenocarcinoma: 637, squamous cell: 223 |
| 5 | TCGA-IntraHeterogeneity | H&E-stained high-resolution image patches of adenocarcinoma and squamous cell lung tissues | Discrimination of different subtypes of cancer (adenocarcinoma vs. squamous cell lung tumors) within high-resolution image patches of TCGA images | 2 classes and 1629 images | adenocarcinoma: 845, squamous cell: 784 |
| 6 | TCGA-InterHeterogeneity | H&E-stained whole-slides images of adenocarcinoma and squamous cell lung tissues | Discrimination of different subtypes of cancer (adenocarcinoma vs. squamous cell lung tumors) within whole-slide images of TCGA images | 2 classes and 1520 images | adenocarcinoma: 761, squamous cell: 759 |
| 7 | BladderScores | IHC-stained images with various staining scores comprising Score 0, Score 1, Score 2, and Score 3 in bladder cancer | Discrimination of different staining scores (Score 0, Score 1, Score 2, and Score 3) of biomarkers | 4 classes and 2137 images | Score 0: 680, Score 1: 235, Score 2: 284, Score 3: 938 |
| 8 | BreastScores | IHC-stained images with various staining scores including Score 0, Score 1, Score 2, and Score 3 in breast cancer | Discrimination of different staining scores (Score 0, Score 1, Score 2, and Score 3) of biomarkers | 4 classes and 2543 images | Score 0: 1817, Score 1: 263, Score 2: 184, Score 3: 279 |

The markers are widely used in clinical immunohistochemistry as biomarkers for detection of various neoplasm types [32, 80]. Several studies have acquired the expressions of biomarkers in biopsy samples of various cancer types to improve the distinction of specific pathological subtyping and understanding of molecular pathways of different cancers. For example, we can refer to the attempts made to discriminate morphologic subtyping of non-small call lung carcinoma (NSCLC), lung adenocarcinoma (LUAD) versus lung squamous cancer (LUSC) [71, 39, 15, 20]. Antiserums staining tissue are sub-classified according to the staining grade. Each tissue sample in this cohort was scored by a trained pathologist using a discrete scoring system (0, 1, 2, 3). A score zero represents no significant protein expression (negative) because there is no staining color, whereas a score three indicates high expression. Positive results were scored based on both the extent and the intensity of staining. For score three, intense staining was required in more than 50 percent of the cells. Other scores including one and two staining comprise in fewer than 50 percent of the total cells [32].

We also obtained the TCGA [62, 61] images by extracting them from the Cancer Digital Slide Archive (CDSA) [27] that is accessible to the public and, at the time of writing this, hosts 31999 whole-slide images from 32 cancer types. For the purpose of this study, we analyze 1520 H&E stained whole-slide histopathology images as well as 1629 H&E stained high resolution image patches (40X magnification) of two TCGA lung cancer subtypes (i.e., LUAD versus LUSC).

### 2.2 Classification and diagnostic framework

This study presents a framework (see Figure 1) to discriminate different cancer types, subtypes, immunohistochemistry markers, and marker staining scores of histopathology images (Table 1). For the first step of our study, the stained whole-slide images with 1504 *×* 1440 and 2092 *×* 975 pixels were obtained from TMA and TCGA databases, respectively. Note that we did not use any pre-processing methods such as color deconvolution to separate the images from staining [79] or any watershed algorithms to identify cells [81] manually. The whole images directly used as the input to the pipeline.

**Figure 1:**
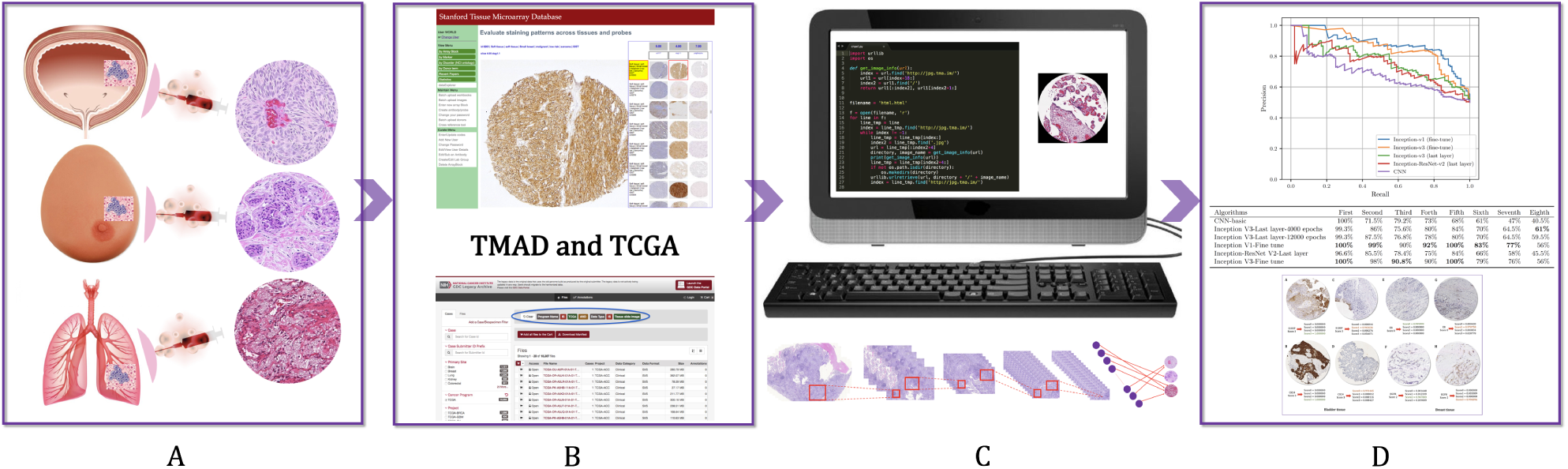
This flowchart demonstrates the pipeline, which includes extracting data, training and evaluation of CNN algorithms, and prediction of various classes. A: tumor image preparation of biopsy samples, B: extracting biopsy-derived tissue slides from TMA and TCGA databases, C: analysis of images using CNN_smoothie, and D: evaluation of the algorithms performance and annotation of the output results.

The images are then divided in different classes based on the classification aims and the CNN algorithms are applied on these classes. For each class, images divided in three groups including training, validation, and test groups. For this purpose, 70% of all images are allocated to the training group and 30% of the remaining images devoted to validation and test sets.

### 2.3 Convolutional Neural Networks (CNNs)

In this study, we use various architectures of CNN algorithms (i.e., deep neural network methods). Neural networks which are the basis of most deep learning approaches comprise certain parameters θ= {*W, B*}, where *W* is a set of *weights* and *B* a set of *biases*. A neural network also consists of neurons with the activation α which represents a linear combination of the input *x* to the neuron and the parameters. In addition, the neural networks contain an element-wise non-linearity *sigma*(.) that refer to a sigmoid and hyperbolic tangent function as α = σ(*w*^*T*^ *x* + *b*). Consequently, the most well-known traditional neural networks is called the multi-layered perceptrons (MLP) that have many layers of transformations. A neural network which contains multiple hidden layers, in between the input and output, is considered a "deep neural network". A survey on deep neural network approaches and their application in medical image analysis is described in [49].

Convolutional neural networks have become the technique of choice for using deep learning approaches in medical images analysis since the first time in 1995 by [51]. Before deep neural networks (DNN) gained popularity, they were considered hard to train large networks efficiently for a long time. Their popularity indebted to good performance of training DNNs layer by layer in an unsupervised manner (pre-training), followed by supervised fine-tuning of the stacked network. In this project, we are going to utilize DNNs for histopathology image analysis. They are the most successful type of models for image analysis because they comprise multiple layers which transform their input with convolution filters [5, 33, 34].

The general concept of a convolutional network is to obtain simple features with higher resolution, and then return them into more complex features at a coarser resolution [73]. The CNNs use the spatial structure of images to share weights across units and benefit of some parameters to be learned a rotation, translation, and scale invariance. So, each image patch around each image can be extracted and directly used as input to CNNs model. One of the very first successful application of deep CNNs was shaped for hand-written digit recognition in LeNet[46]. Then, various novel techniques were developed for training deep networks through efficient ways. The contribution of Krizhevsky and his colleagues [44] to the ImageNet challenge made a watershed advance in core computing systems. They proposed a new architecture of CNN, AlexNet, that won the mentioned competition in December 2012. Currently, the CNNs with deeper architecture and hierarchical feature representation learning have made dramatic changes in object recognition related problems [69, 44, 75, 74, 12].

Simonyan and Zisserman [74] explored much deeper networks containing 19-layer model which called OxfordNet and won the ImageNet challenge of 2014. Then, Szegedy et al. [75] introduced a 22-layer network named GoogLeNet which later referred to as Inception and made use of so-called inception blocks [48], a module that replaces the mapping defined in the 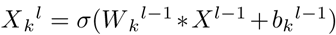 equation with a set of convolutions of different sizes. This Inceptions family architectures allow a similar function to be represented with less parameters. Also, the ResNet architecture [31] won the ImageNet challenge in 2015 and consisted of so-called ResNet-blocks. However, the majority of recent landmark studies in the field of medical imaging use a version of GoogLeNet called Inception-V3 [25, 19, 50]. Recently Esteva et al. [19] utilized a deep CNN as a pixel-wise classifier which is computationally demanding in cancer research to detect melanoma malignant with high performance.

The advantage of Google’s Inception architectures is their good performance even under strict constraints on memory and complexity of computational problems. For example, GoogLeNet [75] used 5 million parameters, which represented a significant reduction in parameters with respect to AlexNet [44] and VGGNet [74]. This is the reason of using Inception networks in big data analysis where huge amount of data needed to be processed at reasonable time and computational cost [59, 72]. Various version of Inceptions are the attempt of Google team to scale up deep networks. For example, in 2014 [75] proposed Inception-V1 and then in 2015 [35] revealed batch normalization. Then, the authors proposed Inception-V2; they presented a derivative form of Inception-v2 which refers to the version in which the fully connected layer of the auxiliary classifier is also-normalized. Then, they call the new model as Inception-v3 which comprising Inception-V2 plus batch-normalization (BN) auxiliary[76]. The Google team also tried various versions of the residual version of Inception such as Inception-ResNetV1 which is high computational cost version of Inception-v3. Another version is Inception-ResNet-V2 that its computational cost matches with the newly introduced Inception-V4 network [77]. However, the Inception-V4 proved to be significantly slower due to the larger number of layers. One of the major technical difference between the residual and non-residual Inception variants is that using BN only on top of the traditional layers in the case of Inception-ResNet [77].

### 2.4 Transfer learning

Image classification was one of the first areas in which deep learning made a principal contribution to medical image analysis. In medical image classification multiple images are considered as inputs with a single diagnostic result as output (e.g., cancerous or normal). A dataset comprising diagnostic image samples have typically bigger sizes with smaller numbers compared to those in computer vision. The popularity of transfer learning for such applications is therefore not surprising that essentially refers a method with two popular and have been widely applied strategies on medical data. Transfer learning refers to pre-train a network architecture on a very large dataset and use the trained model for new classification tasks for a dataset with limited size.

The first strategy includes using a pre-trained network as a feature extractor. A major benefit of this method is not requiring a deep network to be trained and the extracted features smoothly applied to the existing image analysis pipelines [49]. The second strategy is fine-tuning a pre-trained network [49]. Empirical investigation about different strategies have revealed conflicting results. For example, Antony et al. [3] showed that fine-tuning clearly outperformed feature extraction, achieving 57.6 percent accuracy in multi-class grade assessment of knee osteoarthritis versus 53.4 percent. While, [41] showed that using pre-trained network as a feature extractor slightly outperformed fine-tuning in cytopathology image classification (70.5 percent versus 69.1 percent). Besides, two recent published papers presented fine-tuned method by pre-trained version of Google’s Inception-V3 architecture on medical data and achieved a high performance close to human experts [19, 25]. In addition, CNNs developers also train their own network architectures from scratch instead of using pre-trained networks as the third strategy. For instance, Menegola et al. [56] compared few experiments using training from scratch to fine-tuning of pre-trained networks, and indicated that fine-tuning worked better for a small data set (i.e., 1000 images of skin lesions).

Given the prevalence of CNNs in medical image analysis, we focused on the most common architectures and strategies with a preference for far deeper models that have lower memory footprint during inference. In this study, we compare various strategies and architectures for application of CNN algorithm to assess their performance on classification of histopathology images. These are included basic architecture of CNN, pre-trained network (training the last layer) of Google’s Inceptions version 1 and 3, fine-tunning the parameters for all layers of our network derived from the data using two pre-trained version of Google’s Inception architectures (versions 1 and 3), and the ensemble of two the state of the art algorithms (i.e., Inception and ResNet).

### 2.5 Implementation Details

In order to deploy the central architecture, we used a Tensorflow [1] framework. This open source software solution was originally created by the Google Brain team for machine learning applications on textual data sets. The framework supports running the training operation of the network on graphics processing units (GPUs) or traditional computer microprocessors (CPUs). This platform also supports several machine learning algorithms with the same optimizer. The Python programming language version 2.7 was used for all aspects of this project. Also, TF-Slim which is a library for defining, training, and evaluating models in TensorFlow was used in this study. This library enables defining complex networks quickly and concisely while keeping a model’s architecture transparent and its hyperparameters explicit.

A fixed image size of 20 *×* 20 pixels was selected for CNN-basic architecture to ensure that all images have the same size and large cells were entirely captured. CNN with the basic architecture consist of a two layer CNN network with max-pooling blocks; at the end we have two fully connected layers. The image sizes for Inception-V1, Inception-V3, and Inception-ResNet were automatically selected as 224 *×* 224, 229 *×* 229, and 229 *×* 229 pixels by the algorithms, respectively.

All design and training of our method was performed on a desktop computer running the Mac operating system. This computer was powered by an Intel i5 processor at 3.2 GHz, 16 GB 1867 MHz DDR3 of RAM, and a solid state hard drive which allowed ruling out bottlenecks in these components. Although we were able to run all experiments without a GPU (≈7 Gigabyte data), high levels of system memory and a fast storage medium make this application faster since it depends on loading a significant number of medical images for training and validation.

The experimental section is split into two parts: While the aim of the first part of experiment is to reach reliable classification accuracy on the digital pathological images, the goal of the latter is to apply various architectures of CNNs to better understand the choice for the parameters.

### 2.6 Metrics for performance evaluation of algorithms

To assess the performance of different algorithms and to select the most appropriate architectures for a given task and classification aim, we carried out several experiments on the reference datasets. precision-recall curves (PRCs) are typically generated to evaluate the performance of a machine learning algorithm on a given dataset. Recall refers to the fraction of relevant instances that have been retrieved over the total amount of relevant instances, whereas precision measures that fraction of instances classified as positive that are truly positive. In a binary decision problem, a classifier labels either positive or negative can be represented in four categories: true positives (TP) are instances correctly labeled as positives. False positives (FP) refer to negative instances incorrectly labeled as positive. True negatives (TN) correspond to negatives correctly labeled as negative. Finally, false negatives (FN) refer to positive instances incorrectly labeled as negative. Hence, the precision and recall are defined as 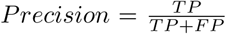 and 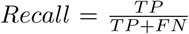. In this study, precisions and recalls are presented by average for multi-class datasets.

To quantify and comparing the performance of various architectures of CNN algorithm on a sample dataset, commonly used accuracy measures, receiver operating characteristic (ROC), were estimated. The ROC curve depicts by plotting the true positive rate (TPR) versus the false positive rate (FPR) at various threshold settings. In ROC plot, FPR locates on the x-axis and TPR on the y-axis. We defined a hard threshold (e.g., from 0 to 1 across a dataset with two classes) for confidence of our predictions. Then, we observe a trade-off between two operating characteristics, TPR and FPR, by varying this threshold. The true positive rate is also known as sensitivity or recall, means the proportion of actual positives in machine learning and false positive rate is also known as (1 *-* specificity) which is the proportion of actual negatives [29, 89]. Therefore, accuracy is measured by the area under the ROC curve (AUC); an area of 1 represents a perfect test and an area of 0.5 shows a worthless test [29, 89].

To evaluate the algorithms performance on all datasets, we also used of two defined measures of accuracy retrieval curve (ARC): true number (TNu) and false number (FNu) [40]. Therefore, to measure the algorithm performances for theses datasets, the accuracy, defined as TNu/(TNu + FNu) which is the fraction of correctly identified images among all images identified by algorithms, while retrieval is the total number of images identified by algorithms.

We also address other measures such as Cohen’s kappa [14] which is a popular way of measuring the accuracy of presence and absence predictions because of its simplicity and its tolerance to zero values in the confusion matrix [2]. The kappa statistic ranges from *-*1 to +1, where +1 indicates perfect agreement and values of zero or less indicate a performance no better than random [14].

The other measure is the Jaccard coefficient measures similarity as the intersection divided by the union of the objects. The Jaccard coefficient ranges between 0 and 1; it is 1 when two objects are identical and 0 when the objects are completely different [13].

The Log-loss or cross entropy which is defined as *-* 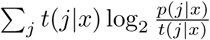 where *p*(*j*|*x*) is the probability estimated by the method for example *x* and class *j*, and *t*(*j*|*x*) is the true probability of class *j* for *x* [8, 18]. It is used to obtain a solution for a wide variety of loss functions and mathematically convenient because it can be computed for each example separately [65, 22, 88].

## 3 Results and Discussions

For the purpose of evaluating our pipeline, we obtained 9649 IHC stained whole-slide images as well as 2490 H&E stained histopathology images of lung, breast, and bladder cancers from TMAD. We also obtained 1520 H&E stained whole-slide histopathology images and 1629 H&E stained high resolution image patches of squamous cell carcinoma and lung adenocarcinoma from TCGA project.In summery, we used eighth different datasets comprising 26 classes (See Table 1). As demonstrated in Table 2, we utilized six state-of-the-art CNN architectures. The first three datasets cover the tasks that are primarily designed for setting up the pipeline (CNN_Smoothie) across different conditions (i.e., discrimination of different cancers and markers). The other datasets refer to challenging problems in clinical context and are designed to assess the application of the pipeline. In addition of investigating different algorithms, we studied the effect of *epoch number* and *training strategies* on the accuracy and compared the performance of various architectures of CNN algorithm for classification and detection of tumor images.

**Table 2:** The results of six state-of-the-art architectures of deep learning algorithms on various datasets using ARC. The numbers show the accuracy percent and are measured based on TNu and FNu. The bold fonts indicate the best classification accuracies on the datasets.

| Algorithms | Tumor type discrimination<br>(bladder, breast, lung) | Bladder biomarker discrimination<br>(GATA3, CK14, S100P, and S0084) | Breast biomarker discrimination<br>(ER, CK17, CK5/6, EGFR, and HER2) | Tumor subtype discrimination<br>in lung (TMAD images) |
| --- | --- | --- | --- | --- |
| CNN-basic | 100% | 71.5% | 79.2% | 73% |
| Inception V3-Last layer-4000 epochs | 99.3% | 86% | 75.6% | 80% |
| Inception V3-Last layer-12000 epochs | 99.3% | 87.5% | 76.8% | 78% |
| Inception V1-Fine tune | **100%** | **99%** | 90% | **92%** |
| Inception-ResNet V2-Last layer | 96.6% | 85.5% | 78.4% | 75% |
| Inception V3-Fine tune | **100%** | 98% | **90.8%** | 90% |
| Algorithms | Tumor subtype discrimination<br>in lung (TCGA intra-<br>images) | Tumor subtype discrimination<br>in lung (TCGA inter-<br>images) | Score discrimination in blad-<br>der (Scores: 0, 1, 2, 3) | Score discrimination in<br>breast (Scores: 0, 1, 2, 3) |
| CNN-basic | 68% | 61% | 47% | 40.5% |
| Inception V3-Last layer-4000 epochs | 84% | 70% | 64.5% | **61%** |
| Inception V3-Last layer-12000 epochs | 80% | 70% | 64.5% | 59.5% |
| Inception V1-Fine tune | **100%** | **83%** | **77%** | 56% |
| Inception-ResNet V2-Last layer | 84% | 66% | 58% | 45.5% |
| Inception V3-Fine tune | **100%** | 79% | 76% | 56% |

### 3.1 Evaluation of various CNN architectures in pathological tumor images

In this section, we present details of our evaluations on various CNN architectures. There are two basic subjects in analysis of digital histopathology images including classification and segmentation [85]. We restricted the evaluations to image-based classification. Also, the basic architecture of CNN was utilized as well as Inception-V1 and Inception-V3 architectures (with fine-tuning the parameters for the last layer as well as all the layers). In addition, we evaluated the ensemble of Inception and ResNet (Inception-ResNet-V2) on all datasets.

Our results show that CNN_Smoothie is able to detect different cancer types, subtypes, and their related markers with highly reliable accuracy which depends on the dataset content, dataset size, and the selected algorithm (Table 2). For example, the pipeline can detect various cancer types by about 100% accuracy (Tumor type discrimination dataset in Table 2). While, the results of cancer subtype detection are varied from 61% to 100% based on the selected database, algorithm architecture, and the presence of heterogeneity in a tumor image (Tumor subtype discrimination datasets in Table 2). In addition, separating various bladder immunohistochemical markers results in 71.5% to 99% accuracy for CNN-basic and Inception-V1 fine-tune, respectively (bladder biomarker discrimination dataset in Table 2). Application of the mentioned algorithms on breast immunohistochemical markers lead to 79.2% and 90% accuracy, respectively (breast biomarker discrimination dataset in Table 2).

Closer look at the Inception-V1 result of bladder cancer (99%) and the related images shows S0084 and S100P were misclassified with GATA3 and S0084, respectively, in two cases out of 200 cases. Moreover, the Inception-V1 result (90%) for discrimination of breast biomarkers revealed that all 10% contradictions have happened between CK17 and CK5/6 due to high similarity between them. This result is in concordant to previous studies such as [78] that compared different IHC markers in breast cancer and showed CK17 and CK5/6 have similar expression patterns.

We configured three datasets (BladderBreastLung, BladderBiomarkers, BreastBiomarkers) to set up the pipeline for all of the proposed experiments. Pathologists typically know what type of cancer each patient has or what marker was used for staining in advance. As expected, the algorithms were successfully able to discriminate various types of cancer with 100% accuracy (Table 2). Furthermore, we designed the experiments to investigate whether keeping the background color might have the potential to introduce certain inherent biases in the datasets and affect the result for discrimination of various markers. The slides across BladderBiomarkers and BreastBiomarkers datasets are stained with different IHC staining colors. However, the results show that Inception architectures (V1 and V3) provide accuracies more than 90% in case of the colored version of the dataset (Table 2). When designing the experiments, we were concerned that the convolutional neural networks might only learn with biases associated to the colors, but the results showed the algorithm’s adaptability in the presence of color information, and their ability to learn higher level of structural patterns typical to particular markers and tumors. This result is in concordance with a previous study that compared three dataset types based on different configurations (i.e. segmented, gray and colored) [57]. Mohanty et al. [57] showed that the performance of the model using segmented images is consistently superior than gray-scaled images, but slightly lower than colored version of the images.

The low concordance of the classification results (by algorithms) for BladderScores and BreastScores datasets (Table 2) to the labels that were determined by pathologists, could be related to the high heterogeneity within tumor cell populations of each slide. Moreover, because we did not have enough images to separate each classes individually, we blended all markers with the same score together (e.g. class score 0 contains GATA3-score 0, CK14-score 0, S100P-score 0, and S0084-score 0). Thus, discrimination of various images in these classes became more challenging. The algorithms are then trained for each score disregard to the markers. Our findings are in agreement with previous studies which showed significant variability between pathologists in score discretization [82, 67, 63, 23, 10, 7, 38] and confirmed that 4% of negative and 18% of positive cases are misclassified even for one type of marker. Consequently, S0084 marker had the minimum cases of misclassification in bladder cancer. Furthermore, the minimum misclassification is related to the score 3 and EGFR marker which is a well known basal marker for breast cancer therapy [45]. Despite the difficulty of the task, the result are comparable with those ones which classified by expert pathologists [80].

Although medical images are mostly interpreted by clinicians, the accuracy of their interpretation is reduced due to subjectivity, large variations across interpreters, and exhaustion [24, 84]. We reviewed BreastScores and BladderScores datasets and the content images that are labeled as negative and positive scores. We found out the low concordance of some our result also could be indeed due to significant human errors in labeling, particularly among positive scores (i.e score 1, 2, or 3) (Figure 2).

**Figure 2:**
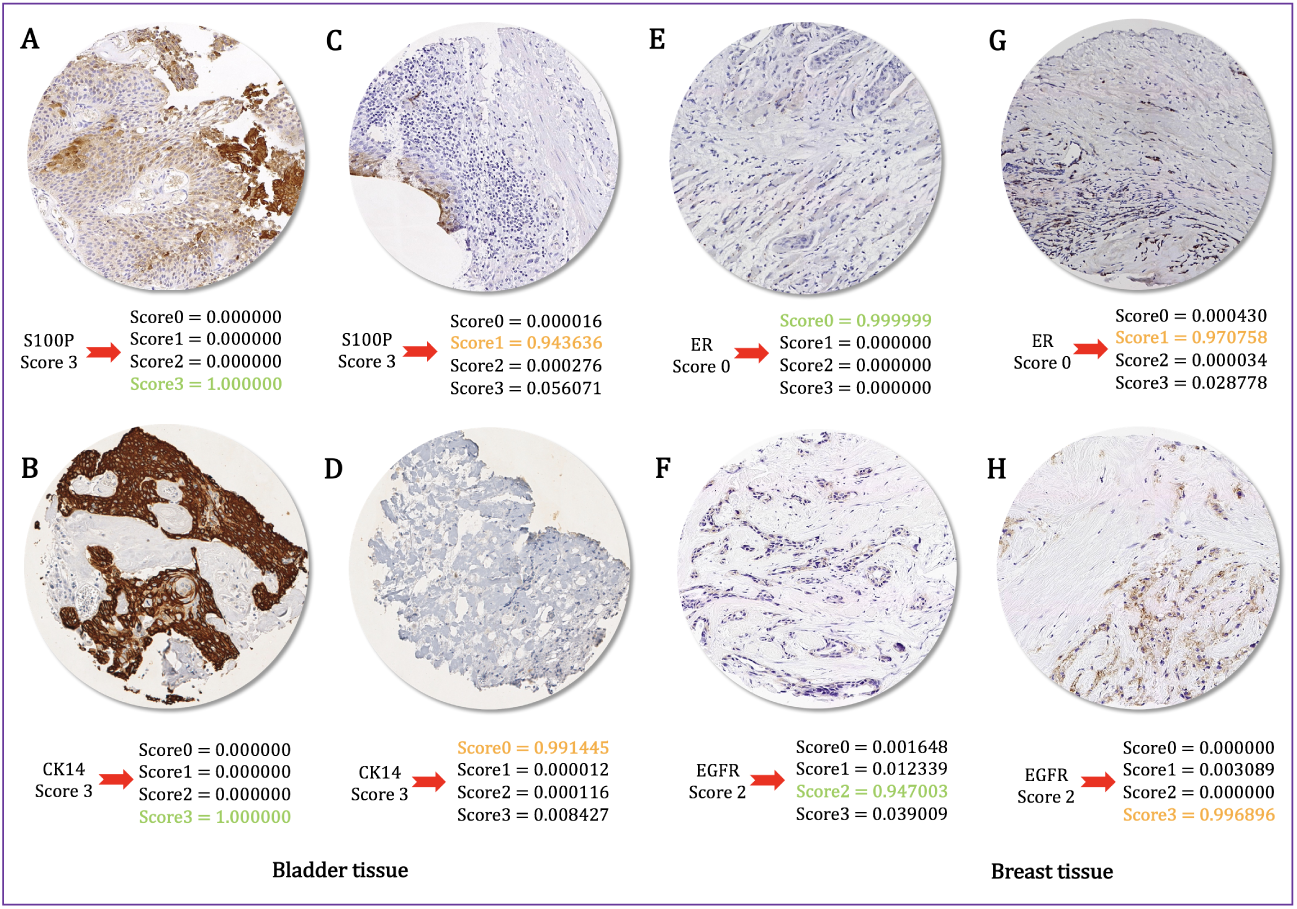
Low accuracy may be related to human errors in labeling the IHC scores. For example, figures A, B, C, and D labeled to score 3 by pathologists, while the algorithm (Inception-V1) has classified them to score 3, 3, 1, and 0, respectively. In particular, figures E and G are both labeled to score 0 by pathologists; however, the algorithm correctly has classified them into score 0 and 1, respectively. Finally, figures F and H are labeled to score 2 by pathologists while the algorithm has classified them into score 2 and 3, respectively. Closer manual inspection of the images indicate the algorithm results are indeed more reliable. Highlighted probability scores in *green* and *orange* indicate concordance and discordance between algorithm classification and pathologist labeling, respectively.

In this regard, we categorized the image datasets into two negative an positive classes for the breast cancer and applied CNN-basic and Inception-V3 (last layer training) on them. The result showed significant increasing of the algorithms performance. The CNN algorithm with basic architecture could discriminate the positive (score 1, 2, and 3) and negative (score 0) images with 94% accuracy. Besides, applying the Inception-V3 which its last layer was trained indicated 96% accuracy for the same dataset.

### 3.2 Discrimination of tumor subtypes across heterogeneous images

Tumor tissues are highly heterogeneous [54] that lead in great limitation for the correct diagnosis. Tumor heterogeneity is the result of genetic disorders which potentially reflects on a variability of morphological features[60].

We randomly selected 1629 H&E stained high resolution image patches (i.e. a few patches of each tumor slide) from TCGA [62, 61] comprising lung adenocarcinoma and squamous cell carcinoma. Then, we trained all CNN architectures for the selected images to discriminate the two subtypes. Consequently, we assessed the performance of the trained algorithms for a separated test set. The test set includes 50 different high resolution image patches of the tumor slides which we trained the algorithms for them (i.e. we considered it as the Intra-tumor test set) (Figure 3). The result showed that while CNN-basic cannot dedicate various cell populations to each subtype, the complex architectures such as Inception-V1 and -V3 can successfully distinguish adenocarcinoma and squamous cell carcinoma across heterogeneous tissue of the tumor slides with no error (TCGA-IntraHeterogeneous dataset in Table 2).

**Figure 3:**
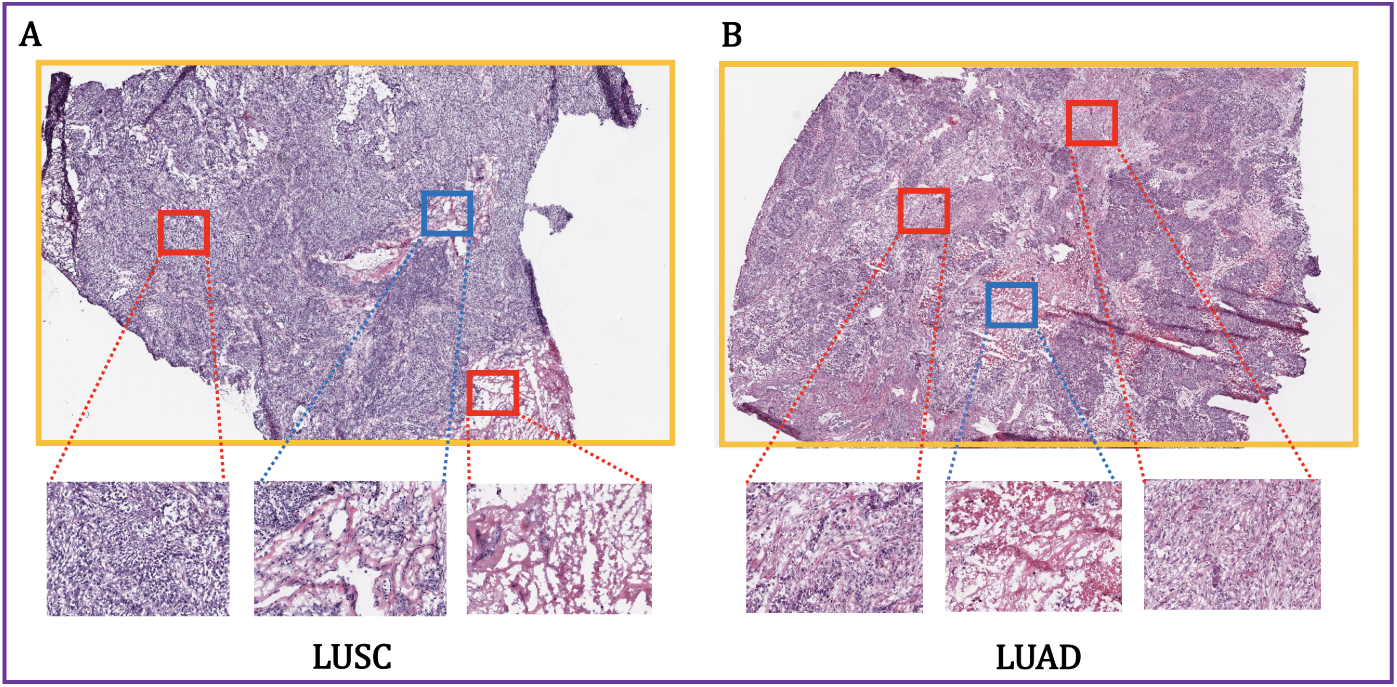
Intra- and Inter-tumor heterogeneity. The figures show the squamous cell lung cancer in the left (A) and adenocarcinoma cell lung cancer in the right side (B). The top images (A and B) represent whole-slide images and the down images represent the extracted high-resolution patches from TCGA datasets. The *red* cubes shows the patches that the algorithms are trained for them and the *blue* cubes indicate the patches comprising test set.

In addition, we assess the performance of the algorithms on inter-tumor heterogeneity of lung cancer. We selected 1520 whole H&E stained histopathology images from TCGA as well as 860 H&E and IHC stained images from TMA database for both lung cancer subtypes (adenocarcinoma and squamous cell carcinoma). Then, we randomly selected and extracted 100 images of each TCGA and TMA datasets separately and trained all algorithms’ architectures for the remaining images. Since the test set images were selected from different patients (tumor slides) that the algorithm never trained for their whole slides or patches, we considered it as Inter-tumor test set. In this way, the algorithms should cope with wide range of cell population variance (intra each individual image and inter different images).

The result indicated 92% and 83% accuracy using the networks which their all layers are fine-tuned based on Inception-V1 parameters for the TMAD and TCGA test sets, respectively (Table 2). The low accuracy of Inter-tumor test set in compare to the Intra-tumor test set can be associated to the high heterogeneity that present across lung cancer for various patients. The mentioned heterogeneity may associated to the various growth patterns (lepidic, acinar, papilary, and solid) [54], grades, and stages in a mixed LUAD and LUSC (or cancer and normal) of the obtained images from various lung cancer patients (Figure 3).

Based on the overall results, it could be useful to use suitable architectures of CNN algorithms based on the goal of the projects. For example, we can use simpler and complex architectures of CNN for discrimination of tumor subtypes through intra- and inter-heterogeneity, respectively. Inter-tumor heterogeneity seems to be more difficult task to detect so needs more complex architecture, while application of the complex architecture on Intra-tumor dataset result in over-fitting and loosing valuable heterogeneity information.

### 3.3 Selecting optimal epoch number and training approach of CNN algorithm

In order to find the optimal epoch number for CNN architectures over different datasets, we stop the training process when the validation accuracy converges to its maximum. We consider that stopping point as the optimal epoch for the tested architecture and dataset (e.g. see Figures 4 and 5). The final classification for images in the test set is performed by re-training the proposed architecture over both training and validation sets with the optimal epoch number.

**Figure 4:**
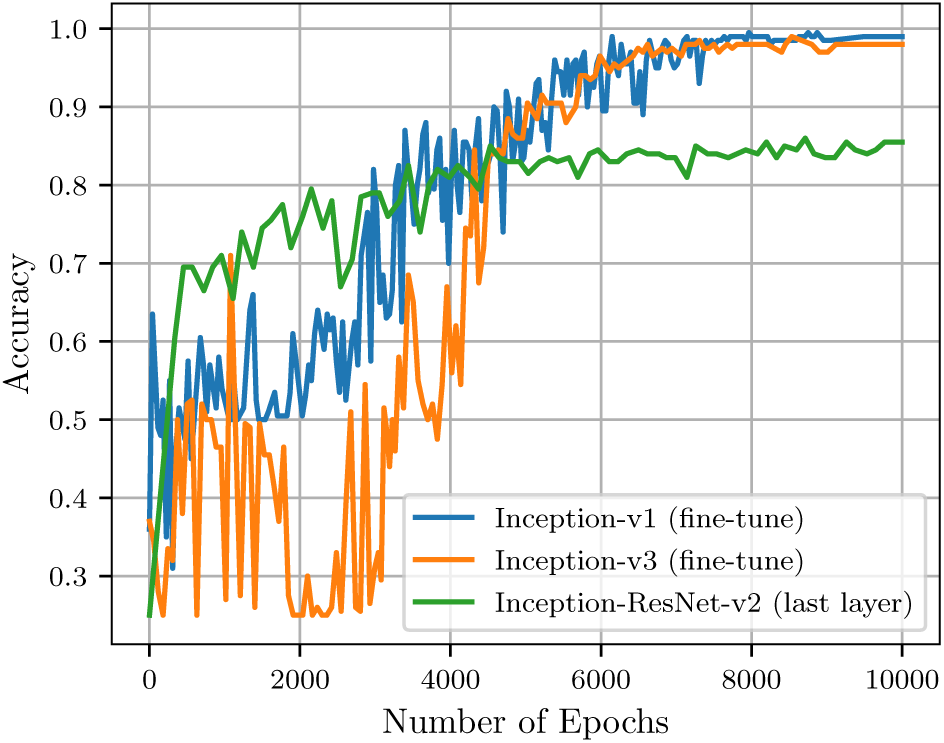
The graph shows the optimal epoch numbers for Inception-ResNet (last layer training), Inception-V1 (fine tuning all layers), and Inception-V3 (fine tuning all layers) to get highest accuracy in BladderBiomarkers.

**Figure 5:**
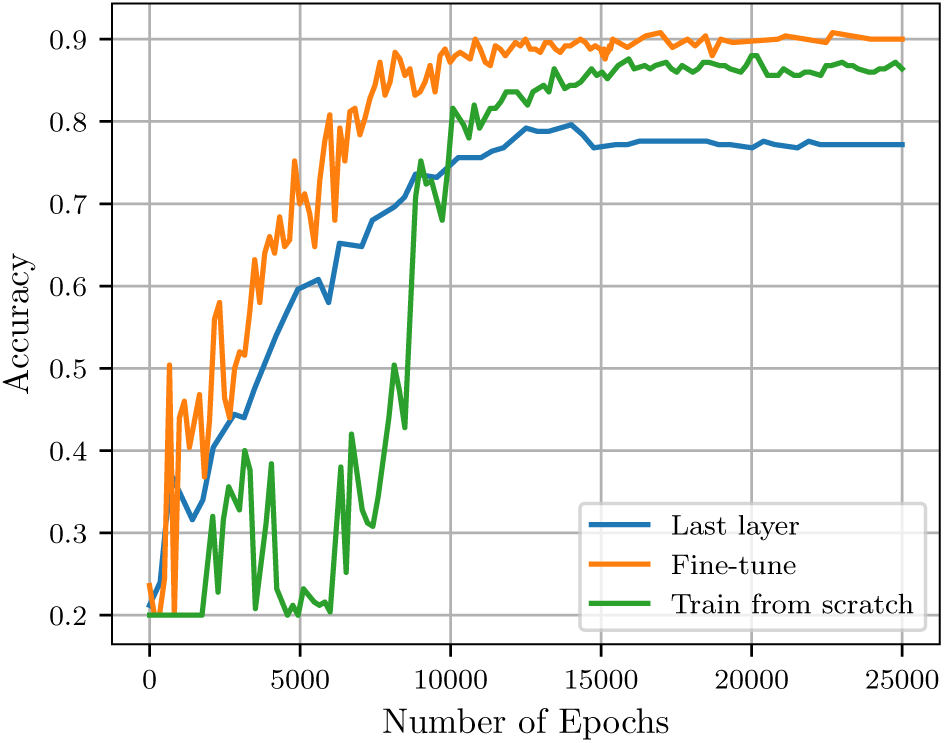
Inception-V1 via three different training strategies (last layer training, fine tuning the parameters for all layers, and training from the scratch) in BreastBiomarkers dataset.

As Table 2 demonstrates, the inceptions-based architecture networks (V1 and V3) that are fine-tuned for all layers, are consistently superior. We also compare various architectures of CNN algorithm using PRC (Figure 6) and ROC (Figure 7 and 8) in one and two sample datasets, respectively, using various thresholds. In this experiment, we consider outputs of an algorithm if prediction’s confidence of a sample pass the determined threshold. We observe a trade-off between precision and recall (for PRC) and TPR and FPR (for ROC) by varying this threshold. These figures reveal that algorithms are able to classify more images which results in a larger recall via smaller threshold.

**Figure 6:**
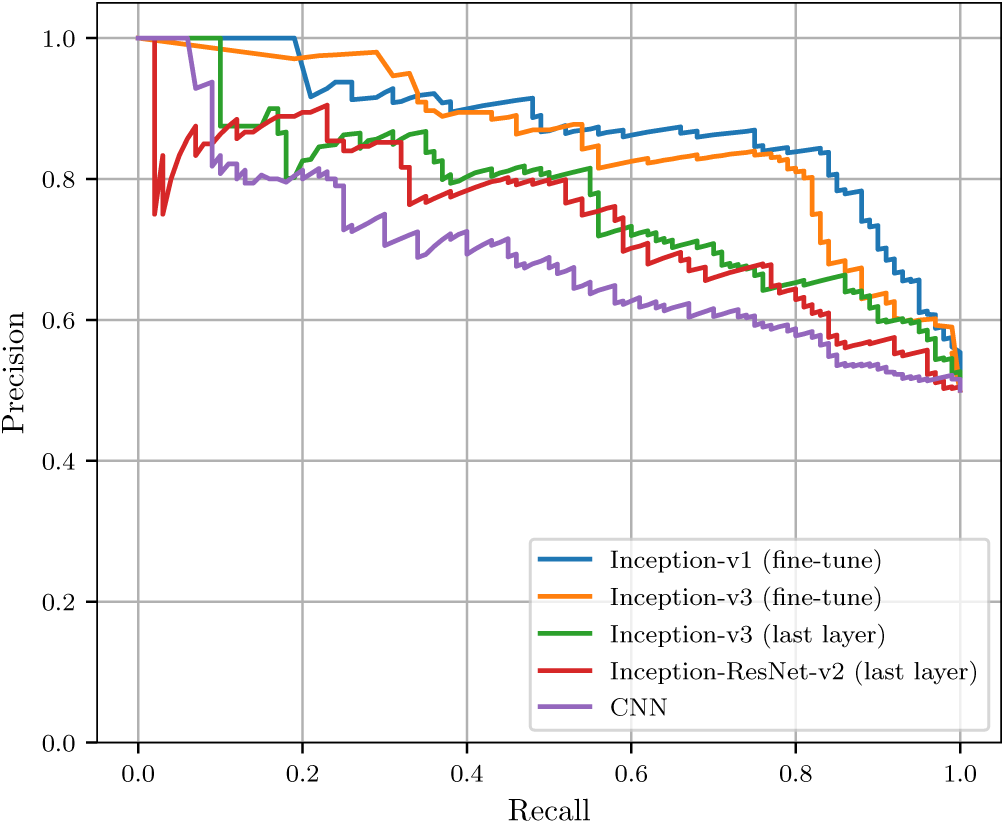
Precision versus recall for the TCGA-InterHeterogeneity dataset. The 4000 epoch version is used for Inception-V3 (training the last layer).

**Figure 7:**
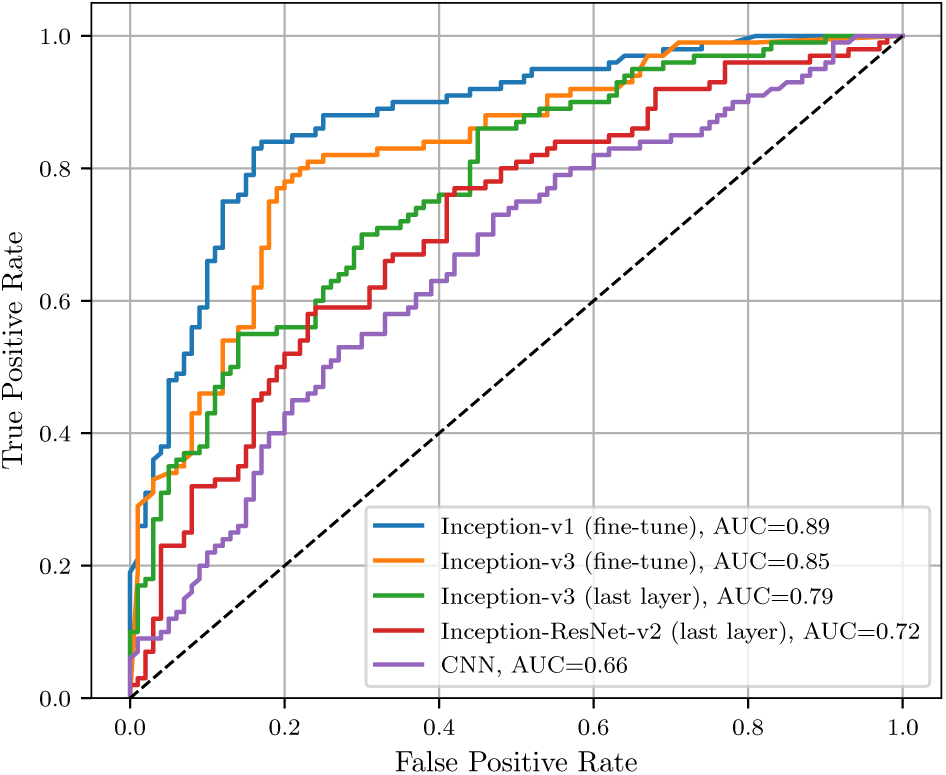
Receiver operating characteristic (ROC) curve for the TCGA-InterHeterogeneity dataset.

**Figure 8:**
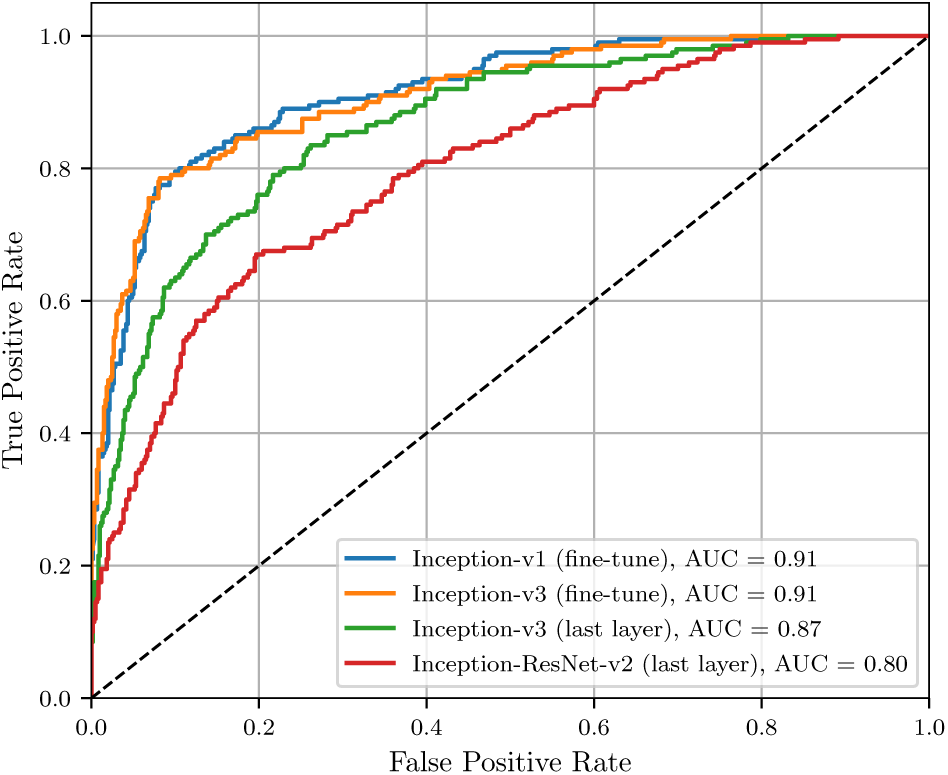
Receiver operating characteristic (ROC) curve for the BladderScores dataset.

We also compare accuracy of different strategies for training Inception-V1. In this regard, we train the model on the marker dataset of breast cancer across training the last layer, fine-tuning of the parameters for all layers, and the training of our own network from scratch (Figure 5). As the figure shows, the best performance is obtained using a pre-trained network and fine-tuning the parameters for all layers of the network, which is in concordance with the results of previous studies [19, 25].

### 3.4 Robustness and limitations of CNN_Smoothie

To demonstrate the robustness of the CNN_Smoothie method, we apply it to eight different datasets of histopathological images with different spectrum of apparent colors to show the uniformity of its performance.The image set spans multiple tumor types, along with several different image colors. The results show that although the colors space for different images have different distributions, our CNN_Smoothie method can successfully identify and register tumor variations and discriminate them consistently and robustly (Figure 9).

**Figure 9:**
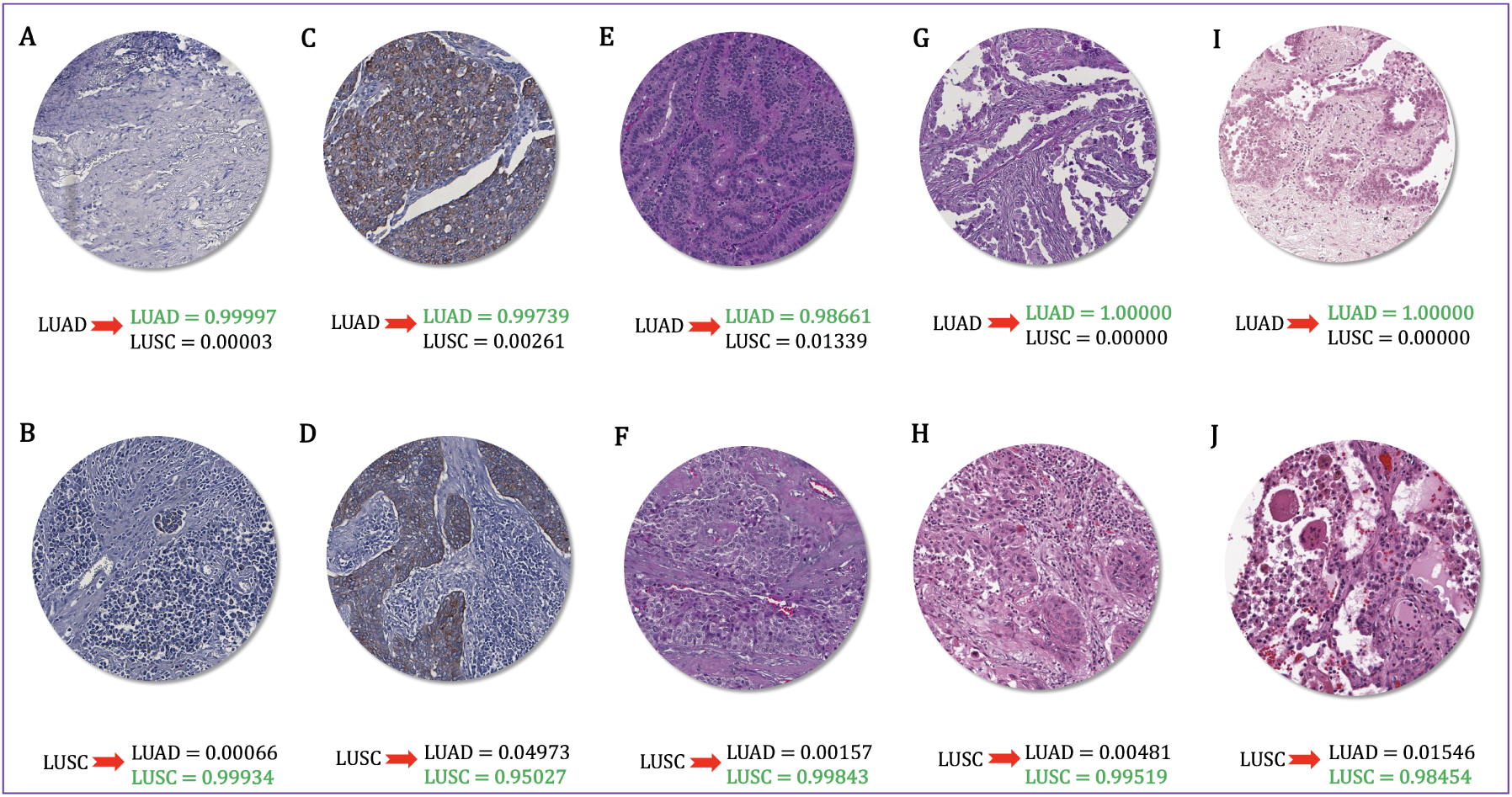
CNN_Smoothie successfully identifies tumor subtypes (LUAD vs. LUSC) and discriminates them consistently and robustly across different spectrum of colors. Highlighted probability scores in *green* indicate the output of classification using Inception-V1.

In addition, we evaluate the performance of algorithms using various statistical measurements on TMAD-InterHeterogeneity and TCGA-InterHeterogeneity datasets to assess the robustness of the results (Table 3). These measures include AUC, average of Precision and Recall, Cohen’s kappa, Jaccard Coefficient, and Log-loss. The Youden index [86] also referred to the ROC which is an indicator for the performance of a classifier and measured as specificity + sensitivity *-* 1 (Table 3).

**Table 3:** The result on TMAD-InterHeterogeneity and TCGA-InterHeterogeneity datasets using various statistics measures. The number in parentheses correspond to the Youden Index. The bold fonts indicate the best classification results for the measures.

| Algorithms | AUC |  | Precision |  | Recall |  | Cohen's kappa |  | Jaccard Coefficient |  | Log-loss |  |
| --- | --- | --- | --- | --- | --- | --- | --- | --- | --- | --- | --- | --- |
|  | TMA | TCGA | TMA | TCGA | TMA | TCGA | TMA | TCGA | TMA | TCGA | TMA | TCGA |
| CNN-basic | 0.64 (0.27) | 0.61 (0.22) | 0.71 | 0.62 | 0.73 | 0.61 | 0.30 | 0.22 | 0.73 | 0.61 | 1.34 | 1.4 |
| Inception-V3 Last-layer 4000-epochs | 0.79 (0.59) | 0.70 (0.40) | 0.81 | 0.71 | 0.80 | 0.70 | 0.56 | 0.4 | 0.80 | 0.70 | 0.45 | **0.57** |
| Inception-V3 Last-layer 12000-epochs | 0.76 (0.52) | 0.70 (0.40) | 0.79 | 0.70 | 0.78 | 0.70 | 0.50 | 0.4 | 0.78 | 0.70 | 0.55 | 0.64 |
| Inception-V1 Fine-tune | **0.89 (0.80)** | **0.83 (0.66)** | **0.92** | **0.84** | **0.92** | **0.83** | **0.81** | **0.66** | **0.92** | **0.83** | 0.39 | 0.66 |
| Inception-ResNet-V2 Last-layer | 0.68 (0.35) | 0.66 (0.32) | 0.74 | 0.68 | 0.75 | 0.66 | 0.38 | 0.32 | 0.75 | 0.66 | 0.48 | 0.63 |
| Inception-V3 Fine-tune | 0.87 (0.75) | 0.79 (0.58) | 0.90 | 0.83 | 0.90 | 0.79 | 0.76 | 0.58 | 0.90 | 0.79 | **0.36** | 1.16 |

## 4 Conclusion

The era of computational pathology is rapidly evolving and there are enormous opportunities for computational approaches to provide additional prognostic and diagnostic information that cannot be provided by pathologists alone [9, 52, 68, 70]. The CNN_Smoothie pipeline presented here provides a novel framework that can be easily implemented for a wide rang of applications, including immunohistochemistry grading and detecting tumor biomarkers. Recently several papers have been published that utilize various methods such as classical machine learning approaches including support vector machine (SVM) and random forest (RF) [87], and deep learning methods such as CNN-basic [80] or Inception methods [19]. However, this is the first report that utilize various architectures of CNN algorithms and compare their performance on histopathological tumor images across various configurations.

The aim of this project is to evaluate the utility of convolutional neural networks to automatically identify cancer cell types, subtypes, related markers, and their staining scores. We indicate deep learning approaches can provide accurate status assessments in clinical conditions. Our results show the accuracy of convolutional neural networks primarily depends on the size, complexity, algorithm architecture, and noise of the dataset utilized. We also show that our study raise several important issues regarding tumor heterogeneity since different response of deep learning could be due to genetic heterogeneity. Further studies required in order to clarify the efficiency of the deep learning application in detection of heterogeneity through digital images.

In terms of computation cost, note that we optimized our pipeline so that it can be run on CPUs. However, GPUs are indeed preferable to scale up the method to Pan-Cancer Analysis and accelerate training speed for future work.

The discordance of our findings and pathology results are due to the low number of tumor images. In certain cases, we blended some images to increase the number of images in each class. In particular, the images associated with biomarkers were blended for each score in BreastScores and BladderScores datasets. Then, the algorithms were trained for different scores disregard of the biomarkers associated with bladder and breast cancers. In addition, the number of images in some classes are not balanced which lead to compliance biases. Finally, we did not train all the algorithms from scratch because GPU is necessary for some datasets and architectures due to their higher complexity. We leave this for future work.

Our method yields cutting edge sensitivity on the challenging task of detecting various tumor classes in histopathology slides, reducing the false rate. Note that, our CNN_Smoothie pipeline requires no prior knowledge of an image color space or any parameterizations from the users. It provides pathologists or medical technicians a straightforward platform to use without requiring sophisticated computational knowledge.

